# High-Speed Neural Imaging with Multiplexed Miniature Two-Photon Microscopy (M-MINI2P)

**DOI:** 10.1101/2025.03.04.641573

**Authors:** Zixiao Zhang, Shing-Jiuan Liu, Ben Mattison, Jessie Muir, Christina K. Kim, Weijian Yang

**Author notes:** These authors contributed equally (Zixiao Zhang, Shing-Jiuan Liu, Ben Mattison). Princeton Neuroscience Institute, Omenn-Darling Bioengineering Institute, Princeton, NJ 08540, USA.

## Abstract

Head-mounted miniaturized two-photon microscopes are powerful tools to record neural activity with cellular resolution deep in the mouse brain during unrestrained, free-moving behavior. Two-photon microscopy, however, is traditionally limited in imaging frame rate due to the necessity of raster scanning the laser excitation spot over a large field-of-view (FOV). Here, we present two multiplexed miniature two-photon microscopes (M-MINI2Ps) to increase the imaging frame rate while preserving the spatial resolution. Two different FOVs are imaged simultaneously and then demixed temporally or computationally. We demonstrate large-scale (500×500 µm^2^ FOV) multiplane calcium imaging in visual cortex and prefrontal cortex in freely moving mice for spontaneous activity and auditory stimulus evoked responses. Furthermore, the increased speed of M-MINI2Ps also enables two-photon voltage imaging at 400 Hz over a 380×150 µm^2^ FOV in freely moving mice. M-MINI2Ps have compact footprints and are compatible with the open-source MINI2P. M-MINI2Ps, together with their design principles, allow the capture of faster physiological dynamics and population recordings over a greater volume than currently possible in freely moving mice, and will be a powerful tool in systems neuroscience.

## INTRODUCTION

Optical imaging of neural activity enables cell type and chemical signaling specific investigation into neural circuits and their correlations with animal behavior. Genetically encoded activity indicators, such as calcium indicators [1–3] or voltage indicators [4–10], provide optical contrast of neural activity through fluorescent microscopes [11–14]. For in-vivo imaging, the animal is typically head-fixed below a benchtop microscope. Virtual reality has further enriched the behavior paradigms [15, 16]. While such benchtop settings have enabled many new discoveries in neuroscience, certain behaviors, such as social behavior, three-dimensional navigation and natural sleeping face challenges when studied in the head-fixed environment. Advances in miniaturized head-mounted microscopes have enabled large-scale imaging of neural activity in the mouse brain during unrestrained and freely-moving behavior [17–37]. In particular, single-photon wide-field miniaturized microscopes or miniscopes [17–23], have been widely adopted in systems neuroscience studies for over a decade. However, single photon excitation is often limited in imaging depth through scattering tissue, as well as signal-to-background ratio due to the lack of optical sectioning capability. Multiphoton microscopy overcomes these limits by using a nonlinear excitation scheme with infrared wavelengths and a point-by-point scanning sampling scheme [38–43]. Recently, there has been a surge in the development of two-photon miniscopes (2P miniscopes) [24–34] and three-photon miniscopes (3P miniscopes) [35–37] for neural activity imaging in freely-moving mice, with the imaging quality on par with basic implementations of their benchtop counterparts. While they are able to image deep tissue with high signal-to-background ratio, multiphoton microscopes have limitations in imaging speed as they build the images pixel by pixel through a single focal spot on the sample. Many techniques, such as beam multiplexing, have been deployed to increase the imaging throughput [44–49]. However, they have not yet been demonstrated in multiphoton miniscopes. Today, the imaging speed of 2P miniscopes is limited by the scanner speed, achieving ∼40 Hz to image 256×256 pixels. While this is sufficient for single plane calcium imaging, it falls short in volumetric multi-plane imaging where different axial planes are sequentially scanned and imaged. Furthermore, it is particularly challenging for 2P miniscopes to perform voltage imaging [6, 8–10], which captures the fast physiological dynamics of the neural activity including both sub-threshold and spiking events, but typically requires an imaging speed of hundreds of Hz to 1 kHz.

In this work, we demonstrate two different multiplexed miniaturized two-photon microscopes (M-MINI2Ps) that each use two fibers to deliver the two-photon laser excitation light to simultaneously image different regions of the FOV. The resulting images are demixed computationally or through temporal gating. Compared to the single fiber case, the resulting imaging speed is doubled while preserving spatial resolution. Using these systems, we demonstrate high-speed volumetric multiplane calcium imaging in primary visual cortex (V1) and prefrontal cortex (PFC) of freely-moving mice (500×500 μm^2^ FOV, single plane frame rate up to 81.5 Hz with 256×256 pixels). During an auditory stimulus, we found the same neurons respond differently in head-fixed or freely-moving conditions. Finally, we demonstrated voltage imaging at 400 Hz in mouse V1 over a 380×150 μm^2^ FOV in freely-moving mouse. M-MINI2Ps extend the open-source MINI2P to overcome speed and throughput constraints of conventional designs while preserving high spatial resolution and FOV. M-MINI2Ps and the beam multiplexing design principle enable high-resolution population voltage imaging and high-speed volumetric calcium imaging over a large FOV in freely-moving mice.

## RESULTS

### Optical design of M-MINI2Ps

The M-MINI2Ps were developed based on the open-source MINI2P and extend its imaging speed and throughput (Figure 1A-B). Instead of a single laser focus, we deliver two focal spots onto the brain tissue and scan them across two different FOVs through the MEMS mirror. We implemented two different multiplexing strategies—temporal (TM-MINI2P) and computational (CM-MINI2P) multiplexing. Both schemes share the same optical layout of the miniaturized microscope, but have distinct benchtop optics optimizing each method and unique demultiplexing methods (Figure 1C-D).

**Figure 1.**
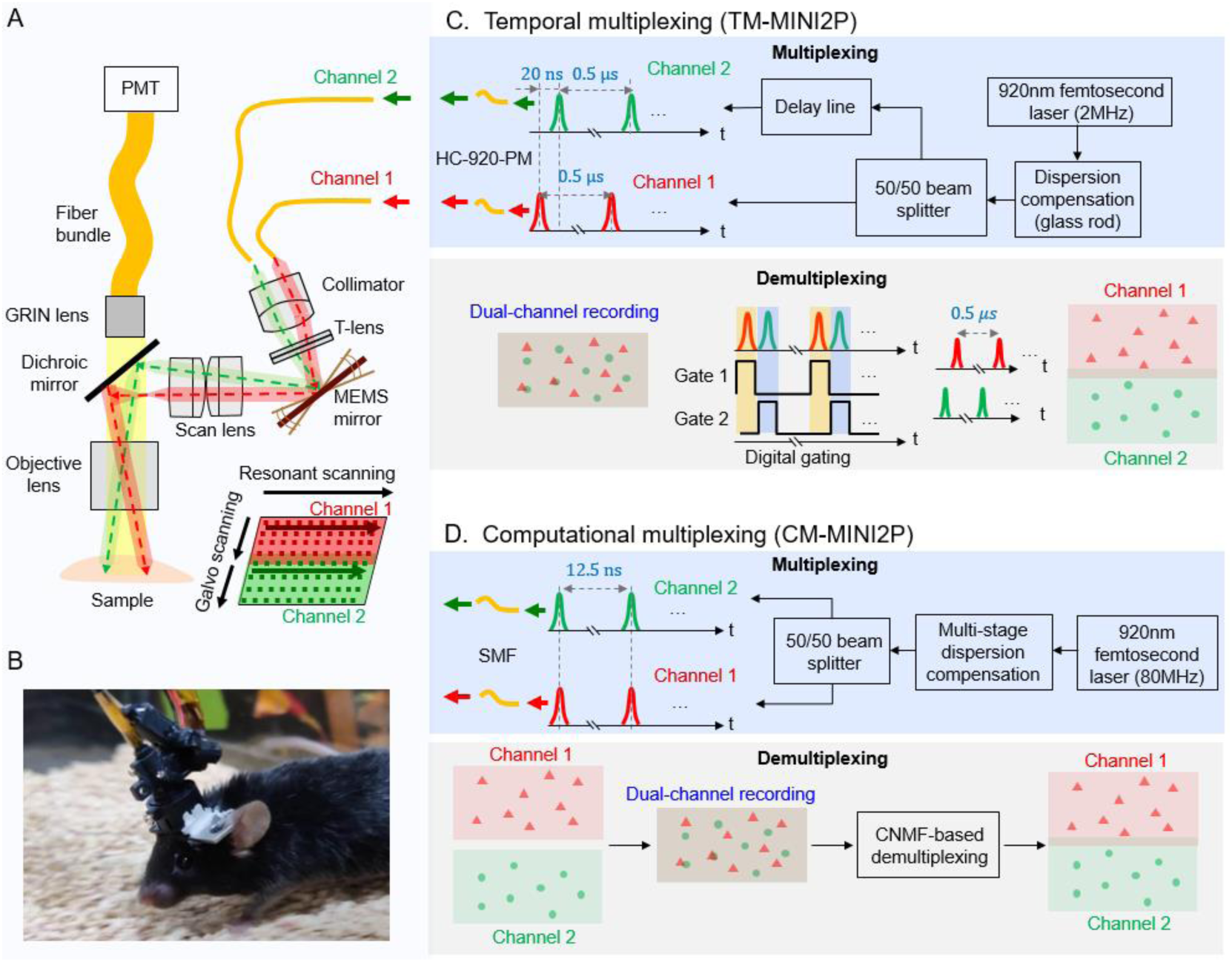
Multiplex miniaturized two-photon microscope (M-MINI2P). (A) Optical layout of the M-MINI2P. Two optical fibers with a lateral displacement deliver separate excitation laser pulse trains to the miniscope. The excitation pathway includes a collimator, a T-lens stack, a MEMS scanner, a scan lens, a dichroic mirror, and an objective lens, enabling spatial multiplexing by directing the laser pulse trains to distinct regions of the sample plane. Emitted fluorescence signals are collected via a fiber bundle and detected by a photomultiplier tube (PMT). (B) Photograph of the head-mounted M-MINI2P attached to a freely-moving mouse. (C) Temporal multiplexing setup (TM-MINI2P). A 920 nm 2 MHz femtosecond laser generates excitation light, which is split into two temporally staggered pulse trains using a 50:50 beam splitter and a delay line. The two beams are coupled into an HC-920-PM fiber, with chromatic dispersion pre-compensated by a glass rod. Digital gating with temporally separated gating windows enables accurate demultiplexing of neuronal signals from each channel, facilitating dual-channel neuronal recording. (D) Computational multiplexing setup (CM-MINI2P). A 920 nm 80 MHz femtosecond laser generates excitation light, which is split into two excitation channels using a 50:50 beam splitter. The beams are coupled into single-mode fibers (SMFs) directed towards the miniscope. A multi-stage dispersion compensation module corrects the dispersion and the nonlinear effects introduced by the SMFs. Neuronal signals from the two channels are separated computationally using a constrained nonnegative matrix factorization (CNMF) algorithm.

In the M-MINI2Ps, two optical fibers are positioned with a lateral displacement from the optical axis (±375 µm) to independently deliver femtosecond laser excitation pulse trains to the miniscope. The excitation pathway consists of a collimator, a tunable micro-lens (T-lens) stack, a MEMS scanner, a scan lens, a dichroic mirror, and an objective lens. The two laser beams from the spatially separated fibers are each collimated after the collimator, incident on the MEMS scanner at different angles, and focused on different lateral positions on the slow scanning axis of the sample plane. The MEMS scanner, conjugated to the back aperture of the objective lens, executes resonant scanning along the fast axis and galvo scanning along the slow axis of the sample plane. The frame rate is determined by the resonant scanning frequency and the number of scanning rows in the slow galvo axis. Compared to the single focal spot configuration, each laser foci in the multiplexed configuration scans half of the original FOV, which effectively halves the number of galvo scanning rows per FOV, thereby doubling the imaging speed (Figure 1A). The MEMS scanner operates at a scanning angle of ±5° along the fast resonant axis and ±2.2° along the slow galvo axis, yielding a total FOV of 500×500 μm^2^. The T-lens stack provides dynamic focal plane adjustment for precise z-scanning and refocusing. Fluorescence emission is transmitted through a dichroic mirror before being coupled into a fiber bundle using a gradient index (GRIN) lens and is detected by a photomultiplier tube (PMT) after the fiber bundle.

The M-MINI2P platform weighs less than 3 g and incorporates flexible cables, ensuring minimal interference with the unrestrained movement in mice, as described in [29]. Locomotion trajectories of the mice with and without the miniscope were compared and analyzed, which shows no significant difference in the accumulated distance traveled and median speed during 10-minute sessions (Supplementary Figure S1). These results verified that the M-MINI2P system did not disrupt animal motion.

### Temporal multiplexing strategy (TM-MINI2P)

In the temporal multiplexing scheme [44] (Figure 1C), a temporal delay is introduced between the two pulse trains towards the two FOVs and time gating at data acquisition is used to separate the signal. Here, we utilize a 920-nm femtosecond laser operating at a 2 MHz repetition rate for excitation. The laser beam is split into two temporally staggered pulse trains through a 50:50 beam splitter and a free-space delay line, with one path delayed by τ = 20 ns. This delay well exceeds the fluorescence lifetime of GCaMP, and allows for temporal separation of the signals between the two channels with minimum crosstalk. The two beams are coupled to two separate 1.5 m hollow-core photonic crystal polarization-maintaining fibers and focus on distinct regions of the sample after the miniaturized two-photon microscope. The MEMS mirror scanning frequency in TM-MINI2P was set to 3.2 kHz along the fast axis, imaging 256 pixels per line with a temporal factor of 0.8. This ensures that each pixel receives a single excitation pulse. Digital gating with temporally separated gating windows during data acquisition at the PMT enables accurate demultiplexing of the signals from each channel. We note that the typical 80 MHz repetition rate femtosecond laser could be used here, but further demixing between the two channels may be required as the temporal delay between them is small.

### Computational Multiplexing Strategy (CM-MINI2P)

Computational multiplexing (Figure 1D) does not require a temporal offset between the two pulse trains, but rather it leverages the temporal dynamics of the neuronal activity to separate the two FOVs which appear overlapped during acquisition [49]. We use a constrained nonnegative matrix factorization (CNMF) algorithm [50, 51] with the guided spatial priors of each individual FOV to demix and extract the neuronal activity of individual regions of interest (ROI, i.e. neurons here) in each of the FOVs [49]. CNMF leverages the spatiotemporal sparsity of the neuronal recording and segments the spatial footprints and extracts the temporal activity of the ROIs through matrix factorization, even when there are spatial overlaps among ROIs. Here, prior to a dual-path multiplexed recording, we first record the individual FOVs by blocking the opposite optical path. CNMF is used to extract the spatial footprints of the neurons, background and noise level of each FOV. This information is then used to initialize CNMF when processing simultaneous two-path recordings.

In this scheme, we opted to use an 80 MHz femtosecond laser as the light source, which allows a higher pixel rate than the 2 MHz laser. The system achieves a frame rate of 81.5 fps at a resolution of 256×128 pixels per single-path FOV, with the MEMS scanner operating at 5.3 kHz along the fast axis. With a greatly reduced peak pulse power, we opted to use to single-mode fibers (SMFs) to deliver the light to the miniscope. This may be more cost-effective compared to photonic crystal fibers, though more sophisticated dispersion compensation is required (Supplementary Figure S2, Methods).

### Multiplexing strategies increase the imaging speed while preserving spatial resolution

We verified that the multiplexing strategies preserve the 3D spatial resolution of the single-beam MINI2P by measuring the point spread function (PSF) of the miniscope with the launch fiber positioned along the optical axis (single-beam MINI2P) and laterally displaced (M-MINI2Ps) from the optical axis of the collimator in the miniscope (Supplementary Figure S3). The FWHM of the lateral/axial PSF was measured to be ∼1.37/23.0 µm for the on-axis launch fiber condition and ∼1.34/24.0 µm for the off-axis condition. Overall, this verifies that the spatial resolution of the miniscope is preserved in the multiplexing configuration and confirms the excitation NA of the miniscope to be ∼0.28.

We also validated the lateral displacement of the two laser foci on the sample plane to be ∼230 µm, and each can image an FOV of ∼270×500 μm^2^ (Supplementary Figure S4).

### Multi-depth calcium imaging of mouse V1 using TM-MINI2P

We applied TM-MINI2P to record the spontaneous neuronal activity in the primary visual cortex of GCaMP6f-expressing mice while the mice freely behaved (Figure 2). Compared to the MINI2P using a single laser beam, only half of the MEMS scanning angle range on the slow axis was used, thus doubling the frame rate. This allowed us to perform high-speed multi-plane volumetric imaging. We used the T-lens stack to switch the focal depth of both beams after each acquired frame. We recorded three axial planes in layer 2/3 of mouse V1, each plane being 500×500 µm^2^, with a volume rate of 16.5 Hz.

**Figure 2.**
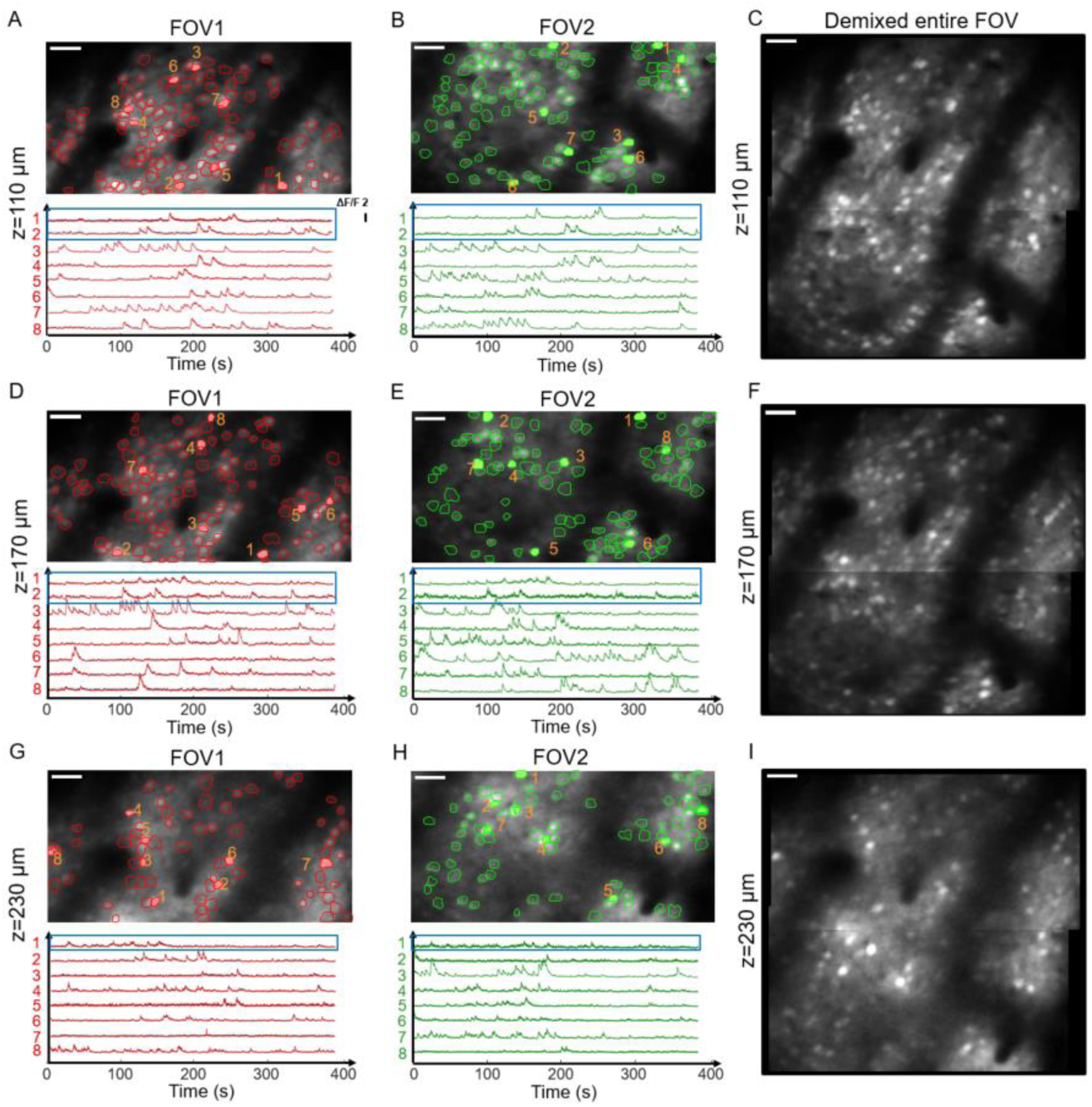
Multi-depth imaging of the spontaneous neuronal activity in mouse V1 using TM-MINI2P while the animal freely behaved. (A-I) Imaging at three depths from pia: (A-C) 110 μm, (D-F) 170 μm, and (G-I) 230 μm in mouse V1 expressing GCaMP6f, with a volume rate of 16.5 Hz. The spatial footprints of the extracted neurons were plotted on the average intensity projections. A total number of 117 (A), 105 (B) neurons were found respectively in FOV 1 and FOV 2 for the 110 μm depth plane, and 119 (D), 96 (E) respectively for 170 μm depth plane, and 72 (G), 70 (H) respectively for 230 μm depth plane. Temporal activity traces for the represented neurons (in shaded colors) were plotted below the average intensity projections. Temporal traces in the blue box represent the neuronal activity of the neurons in the overlapping regions between the two FOVs. (C), (F), (I) The entire FOV stitched from FOV 1 and FOV 2 corresponding to (A-B), (D-E), and (G-H), respectively. Scale bars, (A)-(I), 25 μm.

We adjusted the voltage of the slow axis of the MEMS so that the two FOVs from the two beams had a small spatial overlap. We confirmed that the neurons in the overlap regions can be imaged in both FOVs, with the Pearson correlation coefficient of their temporal traces >0.85 (Figure 2). This further validates the concept of beam multiplexing.

### Multi-depth calcium imaging in mouse V1 using CM-MINI2P

Similar to TM-MINI2P, we validated CM-MINI2P by imaging the neural activity in mouse V1 while the mice freely behaved (Figure 3). We first imaged each individual FOV by blocking the other path. The neuronal footprints were extracted for each FOV by CNMF (Figure 3A-B) and served as the prior information for dual-FOV multiplexed imaging. We then used both paths to perform multiplexed imaging on both FOVs simultaneously. CNMF successfully demultiplexed and extracted the activity traces of the neurons in both FOVs (Figure 3D). Crucially, the average intensity projection of the simultaneous dual-FOV recordings closely matches to the one synthesized by an arithmetical sum of the average intensity projection of the individual FOVs (Figure 3C). Similar to TM-MINI2P, there is a small overlapping region between the two FOVs. Neurons in the overlapping region were imaged twice and appeared twice in the dual-FOV recording (Figure 3D). The Pearson correlation of the activity traces of these repeated neurons (marked with blue masks) was high (>0.9), demonstrating the robustness of the system. By switching the T-lens stack between two focal depths in layer 2/3, we achieved calcium imaging with a volume rate of 40.8 Hz across two depths, each with an effective FOV of 500 × 500 μm² (Figure 3D-E).

**Figure 3.**
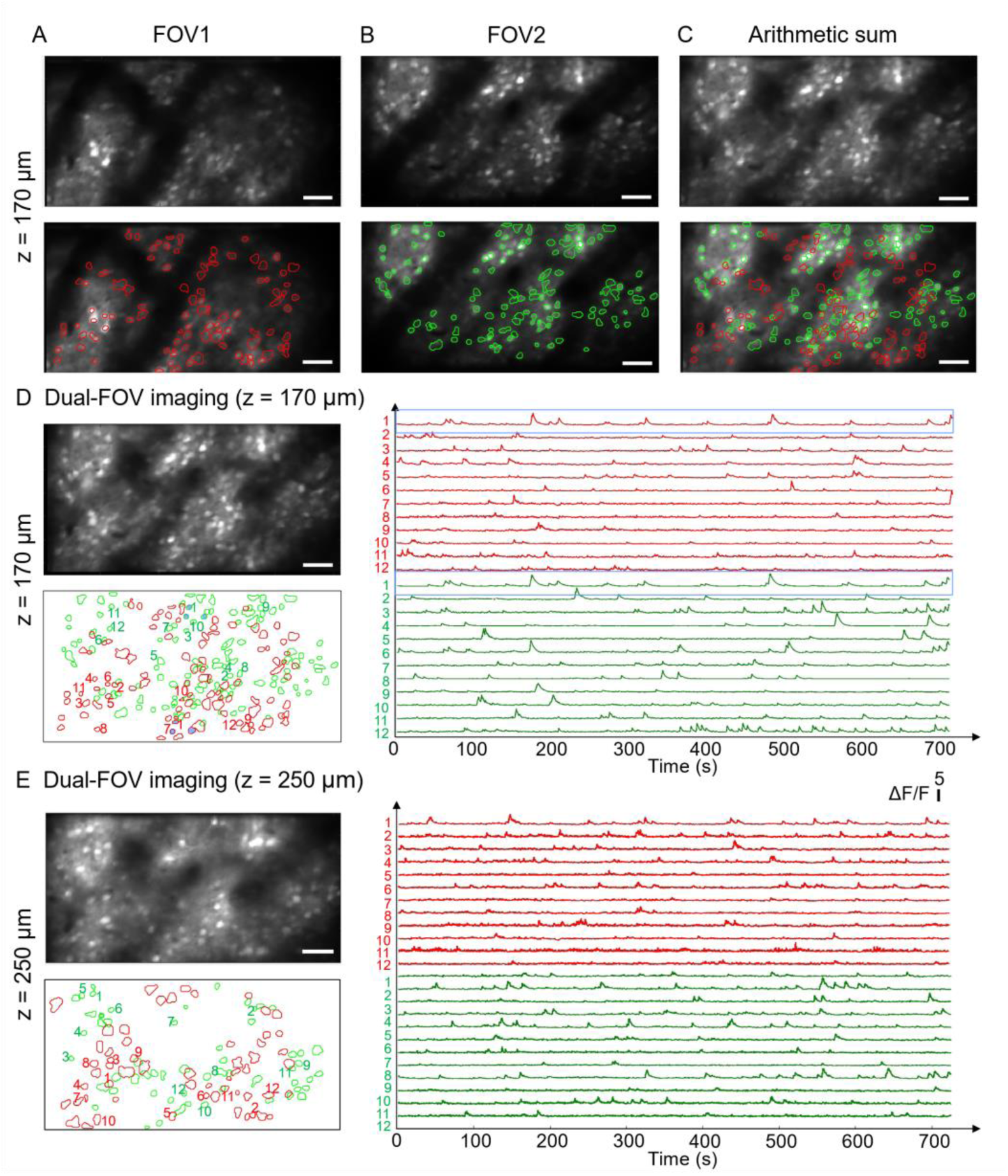
Dual-plane imaging of the spontaneous neuronal activity in mouse V1 using CM-MINI2P while the animal freely behaved. (A) Single FOV imaging of FOV1, with the excitation beam of FOV2 blocked, at the image depth of 170 µm from pia. The spatial footprints of the extracted neurons were plotted on the average intensity projections. (B), Same as (A), but for FOV2, with the excitation beam of FOV1 blocked. (C) Arithmetic sum of the average intensity projections of FOV1 (A) and FOV2 (B), with the superimposed spatial footprints of neurons in both FOV1 and FOV2. (D, E) Dual-depth volumetric imaging at the depth of 170 µm (D) and 250 µm (E) from pia, with a volume rate of 40.8 Hz. Left, average intensity projections (top) and contours of the segmented neurons (bottom). Right, temporal activity traces of represented neurons. The FOV shown in (D) is the simultaneous recording of FOV1 and FOV2 shown in (A) and (B). In (D), neurons in the overlapping regions between the two FOVs are shaped in blue color. The temporal traces of neuron #1 in the overlapping region are shown in the blue box. Scale bar, (A-E), 25 µm.

### Calcium imaging in mouse V1 revealed two distinct social-behavior-related neuronal populations

We demonstrated the capability of our CM-MINI2P system to study the neural dynamics during social behaviors in freely behaving mice, which is challenging in a head-fixed condition. As an example, we examined neuronal activity in mouse V1 during two behavioral conditions: isolated (when the mouse is alone in the cage) and paired (when the mouse could interact with another mouse in the cage) (Supplementary Figure S5). Neurons exhibited distinct activity profiles depending on the behavioral context. We found a subset of neurons that displayed increased neuronal activity during the social interactions, which had more frequent and pronounced calcium transients observed during “paired” sessions, whereas another subset of neurons that were more active during “isolated” sessions. These findings highlight the system’s ability to resolve complex, context-dependent neural activity and establish it as a powerful tool for investigating brain function in naturalistic settings.

### Calcium imaging in PFC during head-fixed and freely-behaving condition

One advantage of using miniscopes is the ability to compare the activity of the same neuronal populations in head-fixed conditions and freely-moving conditions. Previous comparisons have been done using behavioral [52, 53] and electrophysiology data [54]. Two-photon miniscopes are able to provide layer-specific cellular recording comparisons. Here, we investigated if neurons in mouse PFC, which plays a critical role in cognitive functions including fear processing and decision-making, would react differently to auditory stimuli between head-fixed and freely-moving conditions. We performed volumetric calcium imaging in mouse PFC using a GRIN lens implant (Figure 4). The M-MINI2P system enables us to image more neurons through multiple depths in PFC while maintaining a high volume rate (16.5 Hz over 3 planes, with an FOV of 500×500 μm² in each plane). By keeping the M-MINI2P on the mouse’s head, we were able to image the same FOV and volumes in the mouse PFC during head-fixed and freely-moving conditions.

**Figure 4.**
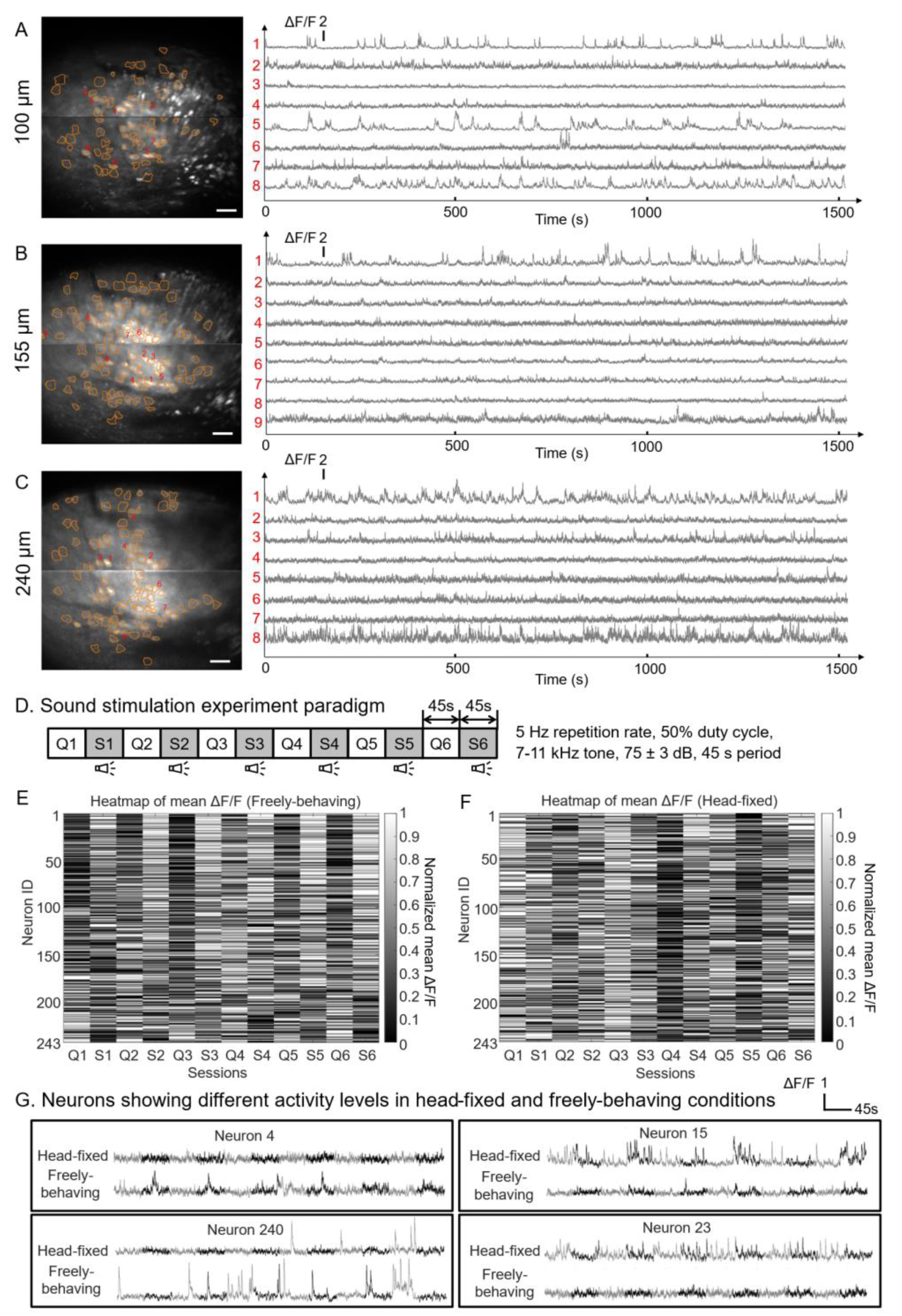
Multi-depth calcium imaging in mouse PFC using M-MINI2P during auditory stimuli. (A-C) Average intensity projections of the calcium imaging at three different imaging depths (100 μm, 155 μm, 240 μm from the end of the implanted GRIN lens) in mouse PFC during freely-moving condition, with the spatial footprint of the extracted neurons overlaid. Temporal activity traces for the represented neurons were plotted on the right. (D) Auditory stimulation experiment with a repeated 5 Hz train (50% duty cycle) of 7–11 kHz tones alternating with quiet periods at 45-second intervals. (E, F) Heatmaps of the normalized mean ΔF/F of all CNMF-extracted neurons under auditory stimulation, recorded in (E) the freely moving condition and (F) the head-fixed condition. Each row represents a neuron, and each column corresponds to a session. The session alternated between sound (S) and quiet (Q) conditions. Neurons were ranked based on the mean ΔF/F between the quiet period and the auditory stimulation period in the freely-behaving condition. Neuron IDs are consistent between (E) and (F). (G) Neuronal activity traces of the exemplary neurons showing different activity levels between head-fixed and freely-behaving conditions (gray: quiet session, black: sound session). Scale bar, (A-C), 25 μm.

We performed auditory stimulation experiments in both freely moving and head-fixed conditions while maintaining the same imaging FOV (Supplementary Figure S6). A repeated 45-second sound-quiet session was used (Figure 4D) for stimulation. To analyze neuronal responses, we extracted neuronal activity traces using CNMF and computed the normalized mean ΔF/F for all detected neurons across sessions. The FOVs in the two conditions were spatially aligned based on maximum spatial correlation, and the same spatial footprints extracted by CNMF were used.

Heatmaps of the activity of PFC neurons revealed distinct response patterns under freely-behaving and head-fixed conditions (Figure 4E-F). In the freely-behaving condition (Figure 4E), we observed clear alternating neuronal firing patterns corresponding to the sound stimulation paradigm. Neurons were classified into three groups: (1) sound-preferred neurons, which were more active during sound sessions (top rows in heatmap); (2) quiet-preferred neurons, which were more active during quiet sessions (bottom rows in heatmap); and (3) non-preferred neurons, which showed no clear preference (middle rows in heatmap). In contrast, this structured response pattern was less evident in the head-fixed condition with the same neuron indices (Figure 4F), and the number of sound-preferred neurons and quite-preferred neurons drops (Supplementary Figure S7). A subset of neurons showed significantly different activity level between freely-behaving condition and head-fixed condition under auditory simulation or quiet condition (Supplementary Figure S7).

In addition to auditory-stimuli-specific responses, we found neurons showed significantly different activity levels between head-fixed and freely-behaving conditions, regardless of the auditory stimuli (Figure 4G). Some neurons exhibited increased activity in the head-fixed condition (Neuron 15 and 23), while others were more active in the freely-behaving condition (Neuron 4 and 240).

These differences suggest that locomotion and behavioral state may influence cognitive functions in the PFC, and that conclusions drawn on head-fixed conditions may not be applied to freely-behaving conditions.

### High-speed voltage imaging in mouse V1

The M-MINI2P system provides a solution for high-speed, high-resolution voltage imaging in freely-behaving animals. Compared to calcium indicators, voltage indicators provide the access to sub-threshold membrane voltage dynamics and fast spiking events. Voltage imaging requires kHz to sub-kHz frame rate. To reach such a high frame rate in two-photon microscopes, the FOV is typically very small. M-MINI2P is advantageous in increasing the FOV while maintaining a high frame rate and spatial resolution. We conducted voltage imaging in mouse V1 expressing ASAP4e [8] over different depths (140 µm, 200 µm, 240 µm, and 280 µm, respectively), with an FOV of 380×150 µm^2^ at each depth, at 400 Hz (Figure 5). We could extract the neurons as well as their spiking events, which were quantified as local peaks exceeding a noise-dependent threshold by a factor of 3.5. By changing the bias voltage of the MEMS mirror at the slow-scanning direction, we could image laterally shifted FOVs in the slow axis at the same depth during sequential sessions. These FOVs could be subsequently stitched into a larger FOV, enabling access to a larger imaging region and allowing investigation of voltage dynamics across wider populations of neurons. To the best of our knowledge, this represents the very first demonstration of high-speed population voltage imaging through two-photon miniaturized microscopes in freely-moving mice. Such an advancement enables precise observation of rapid membrane potential changes, providing critical insights into fast neural computations and circuit-level interactions in freely-behaving animals.

**Figure 5.**
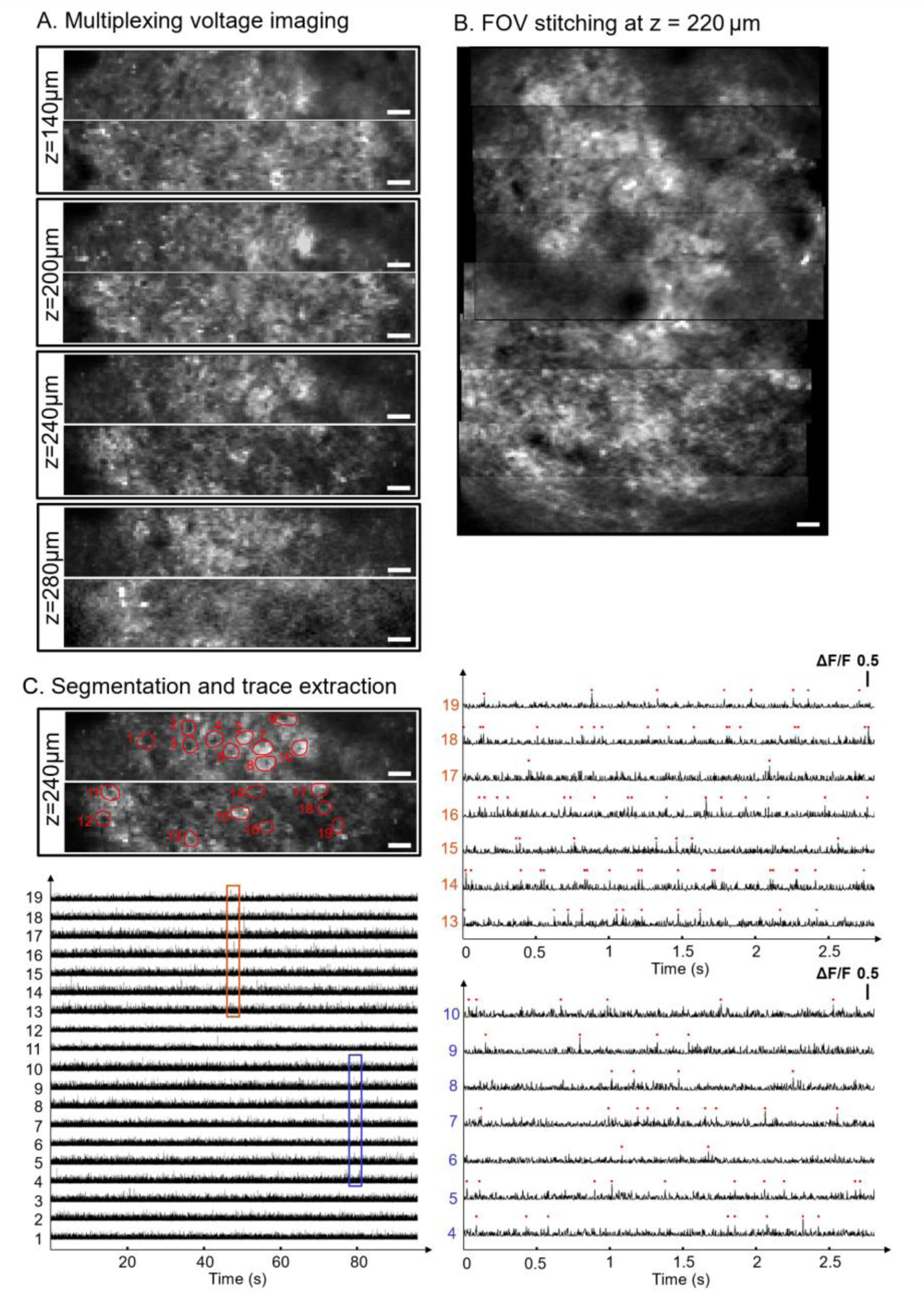
M-MINI2P enables two-photon voltage imaging with a FOV of 380×150 µm^2^ at a 400 Hz frame rate in freely-behaving mice. (A) Average intensity projections of the voltage imaging at different imaging depths from pia (140 µm, 200 µm, 240 µm, and 280 µm) in mouse V1. (B) FOV stitching achieved using MEMS voltage offset. Images at the depth of 220 µm from pia in mouse V1 were stitched to demonstrate access to the full FOV of 380×500 µm^2^. (C) Top: average intensity projections of the voltage imaging at 240 µm imaging depth, with the spatial footprints of the extracted neurons overlaid. Bottom: ΔF/F traces of the extracted neurons. Orange and blue rectangles highlight two distinct time windows, where the traces were zoomed-in in the right panel. Spiking events (red dots) were identified as local peaks rising above a noise-dependent threshold value. Scale bars, (A-C) 25 µm.

## DISCUSSION

We developed two beam multiplexing methods to increase the imaging speed of miniaturized two-photon microscopes and applied them to record neural activity in freely-moving mice. Specifically, we scanned two laser spots on the sample and thus increased the frame rate by 2× compared to the state-of-the-art. This enabled us to perform large-FOV, multi-depth calcium imaging in V1 and PFC with a high volumetric stack rate, and for the first time in miniaturized microscopes, two-photon voltage imaging in freely-moving mice. Our work leverages two advances in multi-photon microscopy: beam-multiplexing techniques in benchtop microscopes [44–49] and head-mounted miniaturized microscopes [24–37]. We engineered the beam launching and collimation module to integrate the beam-multiplexing feature in our miniaturized microscope. Crucially, this does not alter the optical quality of the focal spot on the sample, nor negatively impact the free movement of the mice. While our method is built on the open-source platform MINI2P [29], it can be easily adopted by other miniaturized two-photon or three-photon microscope platforms.

Our beam multiplexing system sets a foundation for further improvement of the optical throughput. More launching fibers could be implemented to increase the number of beams. As the launching fiber has a small diameter (240 µm for PhC and 250 µm for SMF) and is flexible, we expect the miniaturized microscope could accommodate more additional fibers without impeding natural behavior. Further, with the potential of using silicon photomultipliers (SiPMs) for on-board detection as in other 2P and 3P miniaturized microscopes [32, 33, 36], the collection fiber bundle (0.734mm core diameter) could be removed, further untethering the animal. For computational multiplexing, it is also possible to engineer micro-optics to split the beam within the miniaturized microscope without using additional beam launching fibers. Finally, the two multiplexing approaches demonstrated here, can be technically combined with other PSF engineering methods [55] to further increase the imaging throughput as long as they are orthogonal to each other.

The capability of M-MINI2P to perform high-speed two-photon voltage imaging over a large FOV in freely-moving animals represents a substantial leap forward in neural imaging. Existing two-photon miniaturized microscopes are primarily used for calcium imaging, which lacks the temporal resolution to study sub-threshold events and fast neuronal dynamics. The M-MINI2P system, which can image a FOV of 380×150 µm^2^ at 400 Hz frame rate with cellular resolution, addresses this limitation. Such innovations pave the way for transformative research in systems neuroscience.

We were able to perform volumetric imaging of PFC activity through GRIN lenses using M-MINI2P. Interestingly, we found the same neuronal populations could encode different information during head-fixed and freely-moving condition. Such finding has important implications in the study of neural circuits, as the conclusion obtained in head-fixed conditions may not be applicable to a more naturistic state during freely-moving. Further systematic investigation will be required to understand the mechanism underlying these changes of the neuronal representation.

One limitation of GRIN lenses is that they have strong aberrations at the edge of the FOV and away from their focal planes [56]. In our demonstration of volumetric imaging through GRIN lens, we have not corrected for GRIN lens aberrations, thus limiting our cell count slightly. One potential way to mitigate this and increase cell count could be to use aberration correction methods such as microprinted aspherical polymer lenses on top of the GRIN lens [57, 58]. For population imaging, neuropil could be reduced through use of soma-restricted indicators [59].

In summary we have developed M-MINI2Ps with multiplexed architectures, which are able to perform fast population calcium imaging over multiple depths and record voltage dynamics in freely-moving mice. Our development broadens the application of miniaturized two-photon microscopes and also introduces and enables new paradigms for further enhancement of their optical throughput.

## METHODS

### Architecture of the M-MINI2P

We developed two M-MINI2Ps with different beam multiplexing schemes: TM-MINI2P with temporal multiplexing and CM-MINI2P with computational multiplexing. Both TM-MINI2P and CM-MINI2P share the same optical layout of the miniaturized microscope body. The miniscope frame to hold the optical components was fabricated using stereolithography (SLA) 3D printing with BioMed Black Resin (TEAM Lab UC Davis, USA). The miniaturized layout consists of a collimating lens (focal length: 6mm, 63-714, Edmund, NJ, USA) to convert two fiber-emitted diverging beams to collimated beams, a T-lens stack with two stackable T-lens (PIF.15/PAF.19, poLight, Tønsberg, Norway) providing an axial scanning range of 170 µm, a MEMS scanner (A7M10.2-1000AL, Mirrorcle, CA, USA) for lateral galvo-resonant scanning with a tilt angle of 30 degrees relative to the optical axis of the collimation lens, a scan lens (focal length: 5mm, D0166, Domilight, Nanjing, China), a dichroic mirror (MIR-4x5x0.2-HT450-650&HR800-1100, Sunlight Ltd., Fuzhou, China), and an objective lens (D0213, Domilight Ltd., Nanjing, China).

In the emission path, the fluorescence is collected by the objective lens, transmits through the dichroic mirror, relayed via a gradient index (GRIN) lens (64-538, Edmund, NJ, USA), and collected by a supple fiber bundle (Core diameter: 0.74 mm, Fiberoptic Systems, CA, USA) placed at the effective focal length of the GRIN lens. The fiber bundle transmits the collected fluorescence to a photomultiplier tube (PMT, H7422-40, Hamamatsu, Japan) after passing through a bandpass filter (FF03-525/50-25, Semrock, NY, USA). The current signal from the PMT is amplified using a transimpedance amplifier (TIA60, Thorlabs, NJ, USA) and digitized by a high-speed vDAQ (Vidrio Technologies, VA, USA). ScanImage (Version 2022, Vidrio Technologies, VA, USA) was used for data acquisition and hardware control.

Two different femtosecond lasers, in repetition rates of 2 MHz or 80 MHz were used in M-MINI2P. Compared with the 2 MHz rate laser, the 80 MHz rate laser allows a faster pixel rate, but requires higher average laser power on the sample. For TM-MINI2P, we opt for the 2 MHz repetition rate laser (Spirit-NOPA-VISIR, Spectra-Physics, CA, USA) as this allows a larger temporal delay between the two channels to avoid crosstalk. For CM-MINI2P, we opt for the 80 MHz repetition rate laser (Axon 920-2, Coherent Inc., CA, USA) for access to more fast axis pixels.

Two different types of launch fibers can be used in M-MINI2Ps, polarization-maintaining hollow-core photonic crystal fibers (PhC) and single mode fibers (SMF). The PhC fibers are suitable for both 2 MHz and 80 MHz rate lasers, whereas the SMFs are suitable for the 80 MHz rate laser as the high peak power of the 2 MHz rate pulses induces large nonlinearities in the fiber. The key advantage of the HC-PhC is its ability to minimize nonlinear distortion in high-power pulses, as the energy is mainly confined within the air-filled core rather than a dielectric medium. The SMFs have a lower cost than the PhC fibers but more sophisticated dispersion compensation optics are needed. In our demonstration, we used PhC for the TM-MINI2P and SMF for CM-MINI2P.

For the TM-MINI2P system, the 2 MHz repetition rate laser and PhC fibers (HC-920-PM, NKT, Copenhagen, Denmark) are used. The laser-emitted pulse train first passes through 35 cm of H-ZF62 glass rods (GLA-7x500-AR800-1100, Sunlight, Fuzhou, China) to pre-compensate for second order chromatic dispersion. It is then split into two paths by a 50:50 beam splitter (BS017, Thorlabs, NJ, USA). One path is temporally delayed by 20 ns using free-space optics before both paths are coupled into two 1.5 m PhC fibers using achromatic lenses (C240TMD-B, Thorlabs, NJ, USA). The beam path with a longer delay time has a 4f 0.25x beam reducer (AC254-030-B and AC254-120-B, Thorlabs, NJ, USA) in front of the coupling lens to compensate for beam expansion over the longer free-space distance. Demultiplexing is performed by applying 16 ns gating windows in the vDAQ using two channels during data acquisition with ScanImage.

For the CM-MINI2P system, the 80 MHz repetition rate laser and two 1.5 m long SMFs (780HP, Thorlabs, NJ, USA) were used as launch fibers. As the pulse could get distorted in the SMFs, it is critical to correct the distortion and optimize the pulse width. During light propagation in SMF, the pulse width will get broadened due to both linear dispersion and nonlinear effects. The primary linear dispersion arises from large positive group delay dispersion (GDD) introduced by the long SMF. This positive GDD can be compensated using a negative GDD compressor, such as a grating pair compressor, which negatively pre-chirps the pulse before entering the fiber. The dominant nonlinear effect is self-phase modulation (SPM), which becomes significant when the laser power inside the core of the SMF exceeds ∼10 mW for an 80 MHz repetition rate [31]. However, SPM narrows the spectrum of negatively chirped pulses, which increases the pulse width—an effect that cannot be fully compensated by negative GDD compressors alone (Supplementary Figure S2B).

To address both dispersion and SPM, we implemented a multistage dispersion compensation module in the benchtop setup for CM-MINI2P (Supplementary Figure S2). This module consists of a pre-compressor (Compressor 1) within the Axon laser and a grating pair compressor (Compressor 2), both providing negative GDD compensation, with a polarization-maintaining single-mode fiber (PMF) (PM780-HP, Thorlabs, NJ, USA) [25, 60] in between to counteract SPM in the SMF. Compressor 1 slightly pre-chirps the pulses before they enter the PMF, enabling controlled spectral pre-broadening in the PMF by the nonlinear SPM effect. This spectrum broadening was fine-tuned to counterbalance the spectrum narrowing effects in the SMF. Meanwhile, the GDD compressors correct for positive dispersion in both the PMF and SMF, with total compensation dependent on fiber length. By adjusting the compensation ratio of the two compressors, the nonlinear SPM effects in PMF and SMF can be effectively balanced. At the end of the dispersion compensation module, the beam is divided into two paths through a 50:50 beam splitter, and each gets coupled to an SMF through a lens.

With this multistage dispersion compensation, light with an input power of 100 mW into the SMF exhibits an autocorrelation width of <200 fs (Supplementary Figure S2C), measured by an FR-103 autocorrelator (Femtochrome Research, Inc., CA, USA) after the 1.5 m SMF. This corresponds to an estimated pulse width of <150 fs, assuming a hyperbolic secant pulse shape.

To maximize power efficiency, a coupling lens set (Supplementary Figure S2A) consisting of a focusing lens (focal length: 1000 mm, ACT508-1000-B, Thorlabs, NJ, USA), a half-wave plate (HWP, WPH10M-915, Thorlabs, NJ, USA), an adjustable iris (ID25, Thorlabs, NJ, USA), and a collimation lens (F260APC-980, Thorlabs, NJ, USA) were used to optimize coupling into the PMF. The focusing lens corrects the beam divergence over a long propagation distance of 1 m, the HWP optimizes polarization alignment for efficient coupling into the PMF, and the iris fine-tunes the beam size after collimation to match the mode field diameter of the PMF. Additionally, a shifted 4f relay (Thorlabs, NJ, USA) is employed to enhance the coupling efficiency into the SMFs. This 4f system serves as a beam reducer and was intentionally offset to compensate for beam divergence during propagation. The coupling efficiencies were 64% and 72% for the PMF and SMF, respectively. The beam after PMF was expanded by a 3x beam expander (AC254-050 and AC254-150, Thorlabs, NJ, USA) before going to Compressor 2 to reduce average power on the grating pair (49-572, Edmund, NJ, USA). The power efficiency of the grating pair is ∼43%.

### Animal experiments and surgical procedures

Animal preparation and experiments followed protocols in compliance with the NIH Guide for the Care and Use of Laboratory Animals and approved by the Institutional Animal Care and Use Committee at University of California, Davis. Wild-type C57BL/6J mice were used in this study (the Jackson Laboratory) and housed in a 12-hr light/dark cycle in ventilated cages with *ad libitum* access to food and water.

The surgical procedures were virus injection and cranial window implantation for V1 or GRIN lens implantation for PFC. Animals were anesthetized with 2% isoflurane and placed into a stereotaxic apparatus (David Kopf Instruments, USA) with heat pad (Kent Scientific, USA) to maintain adequate body temperature throughout the surgery. Following scalp incision, a microdrill (Foredom, USA) was used to make a hole in the skull over either primary visual cortex (V1) [coordinates 2.4 ± 0.4 mm ML, −3.6 ± 0.4 mm AP, relative to bregma;,−0.350 ± 0.50 mm below dura] or prefrontal cortex (PFC) [0.35 mm ML, 1.9 mm AP, −2.25 mm DV, all relative to bregma]. Virus injection (AAV1-hSyn-GCaMP6f [Addgene] for calcium imaging; AAV9-syn-ASAP4e [Stanford University Viral Vector Core] for voltage imaging) was done using a microsyringe pump (WPI, USA) at speeds of 40 nl/min for V1 or 150 nl/min for PFC to total volumes of 500 nl for V1 and 1000 nl for PFC. Specific dilutions for each virus were 1.5 × 10^12^ GC/mL for GCaMP6f and 8.9 × 10^12^ vg/mL for ASAP4e using a buffer solution comprised of ultra pure sterile water, 1M pH 8.0 Tris, 5M NaCl, and 10% pluronic F-68. A titanium headbar with center hole of 7 mm was installed on each mouse using dental cement (Parkell Metabond, USA) for head-fixation during baseplating. For cranial window implantation over V1, a craniotomy of ∼3.5mm diameter centered over the injection site was performed and a sterile coverslip (Warner Instruments) with thickness 170 µm and diameter 3.5 mm was slowly pressed onto the brain surface. The coverslip was first secured with cyanoacrylate (Vetbond, USA) and the rest of exposed bone was further secured with dental cement. Mice were allowed to wake up and recover and monitored for 10 days post-op. Imaging was done at a minimum of 3 weeks post-op to allow for viral expression.

For GRIN lens implantation into PFC, a 1 mm diameter, 4.38 mm length GRIN lens (1050-006242, Inscopix, USA) was slowly lowered into the injection hole to a depth of 2 mm DV relative to bregma and secured using dental cement. Dental cement was used to cover the rest of the exposed bone. Mice were allowed to wake up and recover and were monitored for 10 days post-op. Imaging was done at a minimum of 6 weeks post-op to allow for viral expression and inflammation reduction.

### Baseplate implantation

Mice were habituated to a custom head-fixation treadmill for a period of 3 days. After habituation and viral expression (a minimum of 3-6 weeks), mice were placed on the treadmill. A multiplexed two-photon miniscope was secured to a plastic baseplate (TEAM Lab UC Davis, USA) using a set screw and attached to a 3-axis stage. The miniscope was angularly aligned to the mouse such that the optical axis of the objective lens was perpendicular to the surface of the cranial window or GRIN lens. Micromanipulation of the 3-axis stage while imaging allowed for finding of a satisfactory FOV. Once a satisfactory FOV was found, the baseplate was secured to the existing metal headplate using dental cement.

Baseplates were designed such that a removable internal spacer was placed between the outer baseplate wall and the miniscope body. Therefore, by adjusting the internal spacer positioning, the positioning of the FOV could be adjusted after securing of the baseplate within a ∼2 mm diameter.

### Imaging sessions

After baseplate implantation, the miniscope could be easily attached and released from the mouse using a set screw. Freely moving recordings were done in 40 cm × 26 cm cages. An IR webcam with integrated IR LEDs (SVPRO 1080P Night Vision USB Camera CMOS OV2710, Shenzhen Ailipu Technology Co., Ltd., China) was secured above the cage to illuminate and record mouse navigation movements during perceived darkness for the mouse. Data for sessions of spontaneous activity recording in V1 were captured for 8-12 minutes at a time for up to 7 sessions per day across 3 days. Data for sessions of recording auditory modulated activity in PFC was acquired in sessions of 9-24 minutes and up to 2 sessions per day. Each session included audio tone profiles alternating with 45 seconds of auditory stimuli and 45 seconds of quietness periods for a total of 6-16 on-off cycles.

### Auditory Stimulation

Audio stimulation was performed by a mobile phone (iPhone 13, Apple, USA) placed 30 cm above the open field behavior arena and calibrated to 75 dB +/-3 dB at the center of the arena. Audio files with 5 Hz sequences at 50% duty of 1 kHz, 4 kHz, 7 kHz, and 11 kHz tone frequencies were generated in MATLAB and downloaded to the mobile phone. Experimental sessions used a 45 s audio on and 45 s audio off paradigm.

### Profiling of the animal movement

We compared the animal’s movement with and without the miniaturized microscope (Supplemental Fig. S1). Six mice with cranial window surgery and baseplate implantation were affixed with a dummy miniscope with optical fibers (two excitation fibers and one collection bundle), electrical cables for MEMS and TLens and weight matched to the experimental scopes. The mice were recorded while freely exploring a 40 cm × 26 cm cage for 10 minutes with and without the dummy miniscope, on sequential days. The videos were analyzed by a trained DeepLabCut model [61], where the movement trajectory was extracted (Supplemental Fig. S1 C.). Briefly, the DeepLabCut model extracted the positions of nose, left ear, right ear, and tail base of each mouse over time during each session, and marker positions with a likelihood <0.5 were rejected. The mouse position was calculated as the geometric center of the left ear, right ear, and nose. Instantaneous speed was calculated as the position change over one frame (∼0.03 s). A comparison of cumulative running distance and median speed of the mice during the sessions found no significant difference between statistics of mice with and without the dummy miniscope (p=0.52 and p=0.49 respectively, N=6, paired t-test) (Supplementary Figure S1).

## DATA AVAILABILITY

Data underlying the results presented in this paper are not publicly available at this time but may be obtained from the authors upon reasonable request.

## ACKNOWLEDGMENTS

We acknowledge support from National Institute of Neurological Disorders and Stroke and National Eye Institute (R01NS118289, WY), National Institute of Biomedical Imaging and Bioengineering (R21EB035306, WY), National Science Foundation (CAREER 1847141, WY), Burroughs Wellcome Fund (Career Award at the Scientific Interface 1015761, WY; and Career Award at the Scientific Interface 1019469, CKK), National Institutes of Mental Health (DP2MH136588, CKK), and the Canadian Institutes of Health Research (post-doctoral training award 202210MFE-491520-297096, JM).

## AUTHOR CONTRIBUTIONS

Conceptualization: WY; Methodology: ZZ, SL, BM, WY; Software: ZZ, SL, BM; Validation: ZZ, SL, BM, WY; Formal Analysis: ZZ, SL, BM; Investigation: ZZ, SL, BM, JM; Resources: WY, CKK; Data curation: ZZ, SL, BM, WY; Writing – Original Draft: ZZ, SL, BM, WY; Writing – Review & Editing: WY, CKK, ZZ, SL, BM, JM; Visualization: ZZ, SL, BM; Supervision: WY, CKK; Project Administration: WY; Funding Acquisition: WY, CKK.

## DECLARATION OF INTERESTS

The authors declare no conflicts of interest.

## Supplemental Information

**Figure S1.**
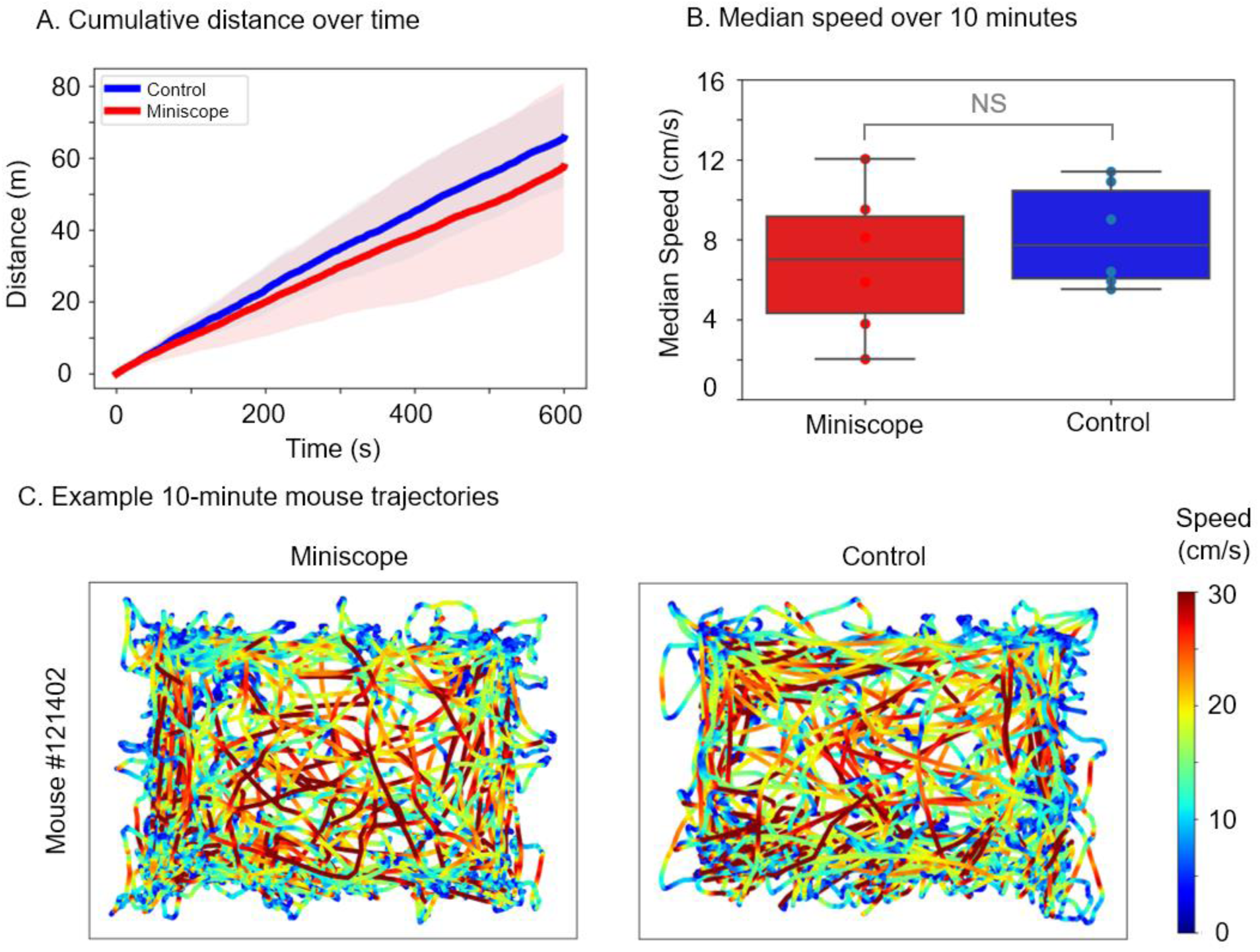
Assessment of free-moving behavior in mice with and without the miniscope system. (A) Cumulative distance traveled during a 10-minute session in a 40 cm × 26 cm open field for mice carrying the miniscope (red line, N = 6) and control condition without the miniscope (blue line, N = 6). Line indicates the mean and shaded areas indicate ± 1 standard deviation. (B) Median running speed of the mice during 10-minute sessions in a 40 cm × 26 cm open field. No significant difference (NS, p=0.49) was observed between the miniscope and control groups (N=6, paired t-test). (C) Representative trajectories during a 10-minute session for a mouse carrying the miniscope (left) and without the miniscope (right). Behavior was tracked by an infrared camera and analyzed with DeepLabCut. Running speed is overlaid and color-coded on the trajectory.

**Figure S2.**
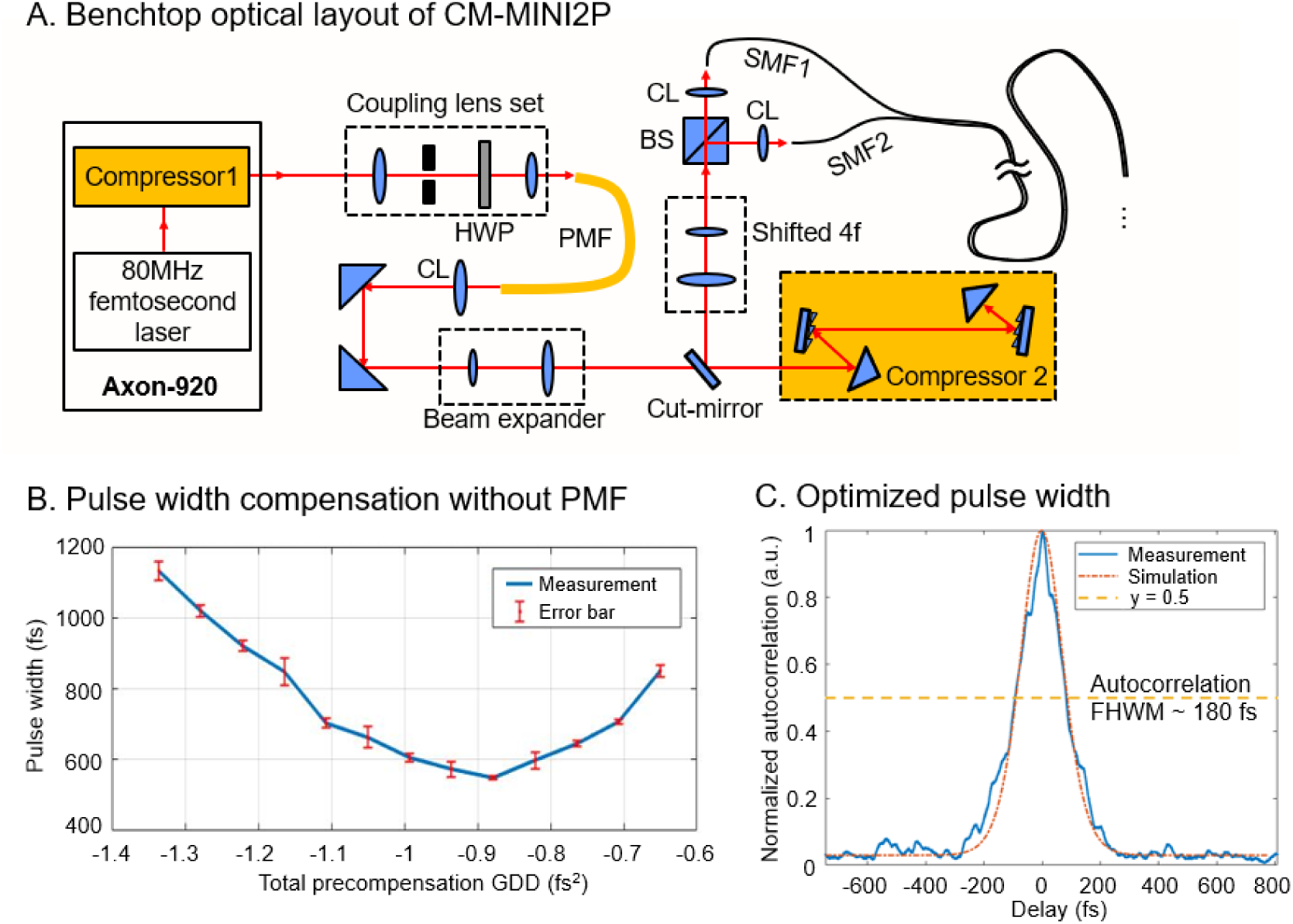
Benchtop optical layout of CM-MINI2P and pulse width optimization. (A) Optical layout of the benchtop components of the CM-MINI2P for pulse width optimization, fiber coupling, and spatial multiplexing. The pulse width of the femtosecond laser is modulated by a multi-stage dispersion compensator, including a pre-compressor (Compressor 1) within the laser and a grating pair compressor (Compressor 2), both providing negative group delay dispersion (GDD) compensation, and a polarization-maintaining fiber (PMF) in between to compensate the nonlinear self-phase modulation (SPM) in single-mode fiber (SMF). A beam expander reduces the average light intensity on the grating pair. The coupling lens set optimizes fiber coupling efficiency into the PMF. The shifted 4f serves as a beam reducer to optimize the beam size and collimation so as to maximize the fiber coupling efficiency into the SMFs. (B) Pulse width measurement at the sample plane without PMF, with 100 mW input power to the single-mode fiber (SMF) with a length of 1.5 m. Nonlinear SPM in the SMF limits the minimum achievable pulse width to >550 fs. (C) Optimized pulse width measurement at the sample plane after the multi-stage dispersion compensation, including PMF, which compensates the SPM in the SMF. With 100 mW input power to the SMF, the measured autocorrelation (averaged over three measurements) of the light pulse has a full width at half maximum (FWHM) of ∼ 180 fs. Assuming a hyperbolic secant pulse shape, the corresponding pulse width is estimated to be ∼110 fs. The simulation curve (red) represents the autocorrelation of a hyperbolic secant pulse with a pulse width of 112 fs.

**Figure S3.**
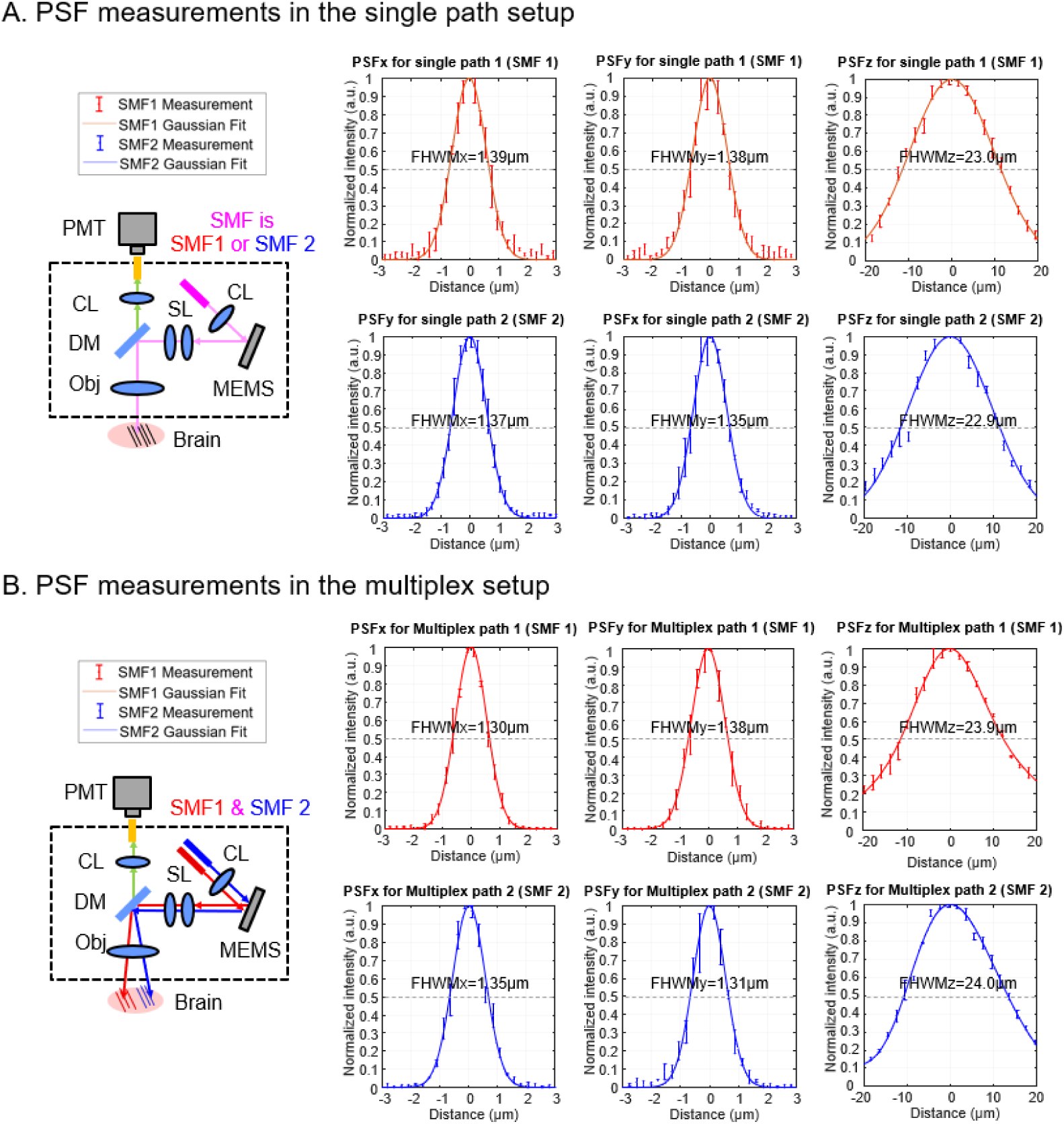
Measurement of the point spread function (PSF). The PSFs were measured with the launch fiber positioned at the optical axis (single-beam MINI2P) and laterally displaced from the optical axis (M-MINI2P) of the collimator at the miniscope, using 500 nm fluorescent beads. The PSF results were averaged with 3-6 beads in each condition, with the standard deviation plotted as error bars. The experiment results were fitted with Gaussian functions, where the FWHM for lateral (PSFx and PSFy) and axial (PSFz) resolutions were obtained. (A) Measurement with a single launch fiber positioned at the optical axis of the collimator at the miniscope. Measurements were performed separately for SMF1 (red) and SMF2 (blue). The lateral PSFs (PSFx and PSFy) ranged from 1.3 to 1.4 µm, and the axial PSFs (PSFz) were ∼23 µm. (B) Measurement with the two launch fibers (SMF1 and SMF2) positioned ±375 µm away from the optical axis. The lateral PSFs (PSFx and PSFy) were ∼1.3 to 1.4 µm, the axial PSFs (PSFz) were ∼24 µm. These results confirm that the spatial resolution was preserved in the multiplex configuration, validating the optical performance of the miniscope under multiplexed operation.

**Figure S4.**
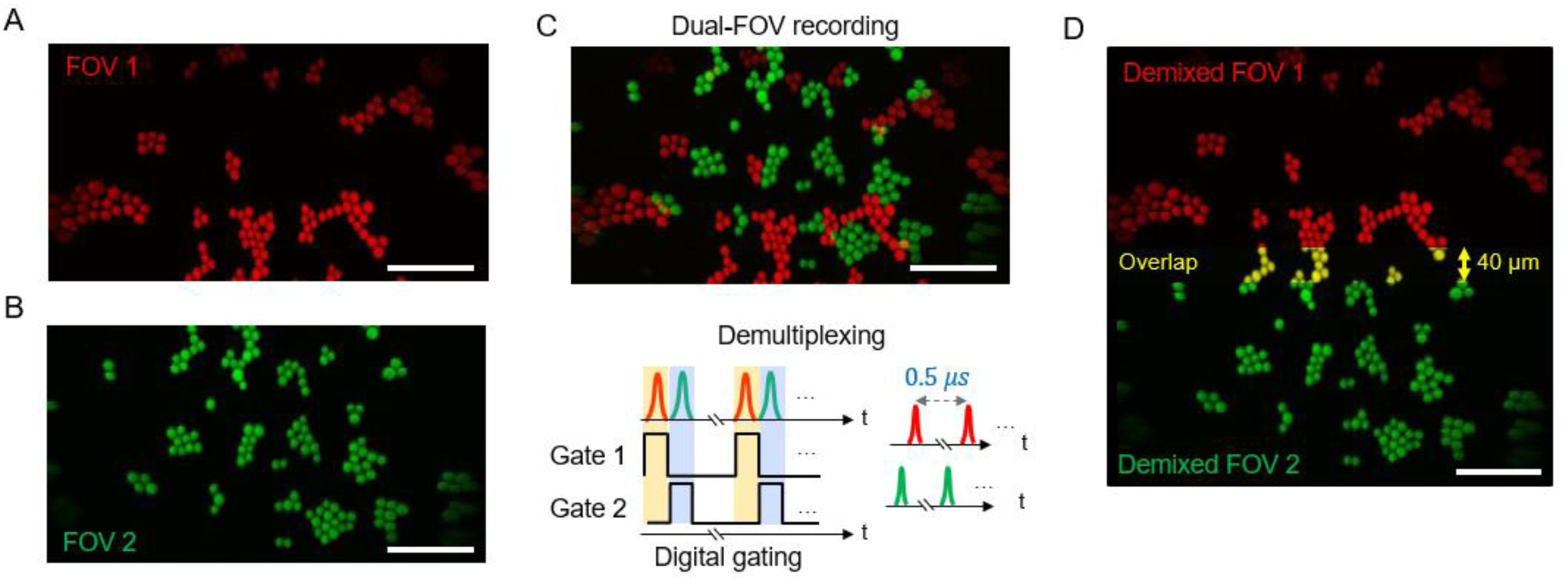
Validation of the temporal multiplexing strategy using a phantom sample with randomly distributed 12 μm fluorescent beads. (A) Pure FOV1 imaging result with the FOV2 excitation beam blocked. (B) Pure FOV2 imaging result with the FOV1 excitation beam blocked. (C) Dual-FOV recording showing signals from both FOVs, separated temporally using digital gating during data acquisition. The temporally demultiplexed signals from each FOV are color-coded accordingly. (D) Reconstructed FOV (500 × 500 μm) after temporal demultiplexing, recovering the original FOVs. The overlapping region (500 × 40 μm) between the two channels is highlighted in yellow, confirming the fidelity of the temporal multiplexing approach. Scale bar: (A-D), 100 µm.

**Figure S5.**
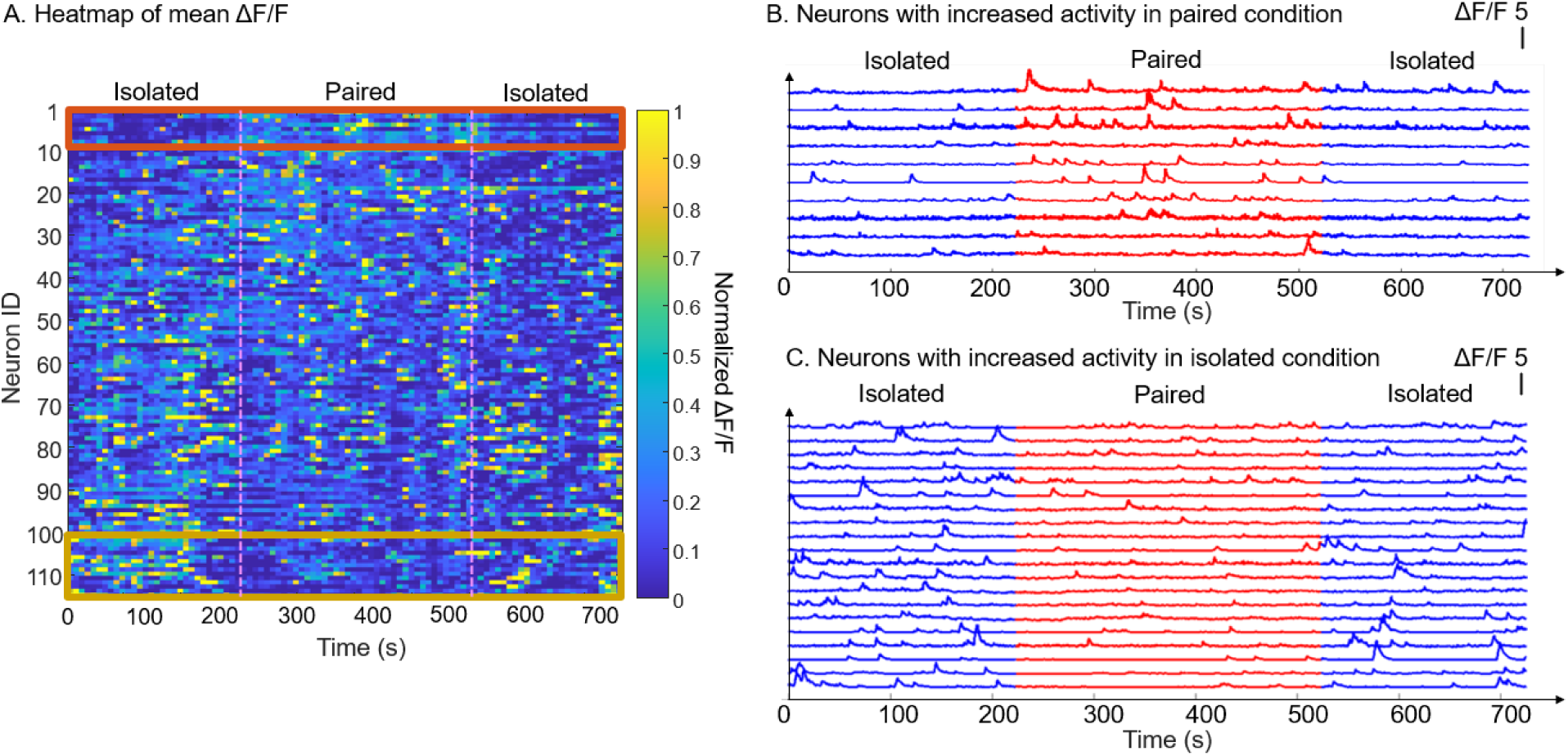
Neuronal activity during social behavior. (A) Heatmap of mean ΔF/F activity for 115 neurons in mouse primary visual cortex (V1) during isolated condition when the mouse was alone in the cage and paired condition when the mouse could interact with another mouse in the cage. The recording was binned to 7.5 sec for visualization. Neurons were ranked by the difference between the mean ΔF/F activity during the paired condition session and the isolated condition session. (B) Activity traces of the representative neurons which exhibited higher activity levels during the paired condition sessions (red), characterized by more frequent calcium transients during social interactions. (C) Activity traces of the representative neurons which exhibited higher activity levels during isolated condition sessions (blue).

**Figure S6.**
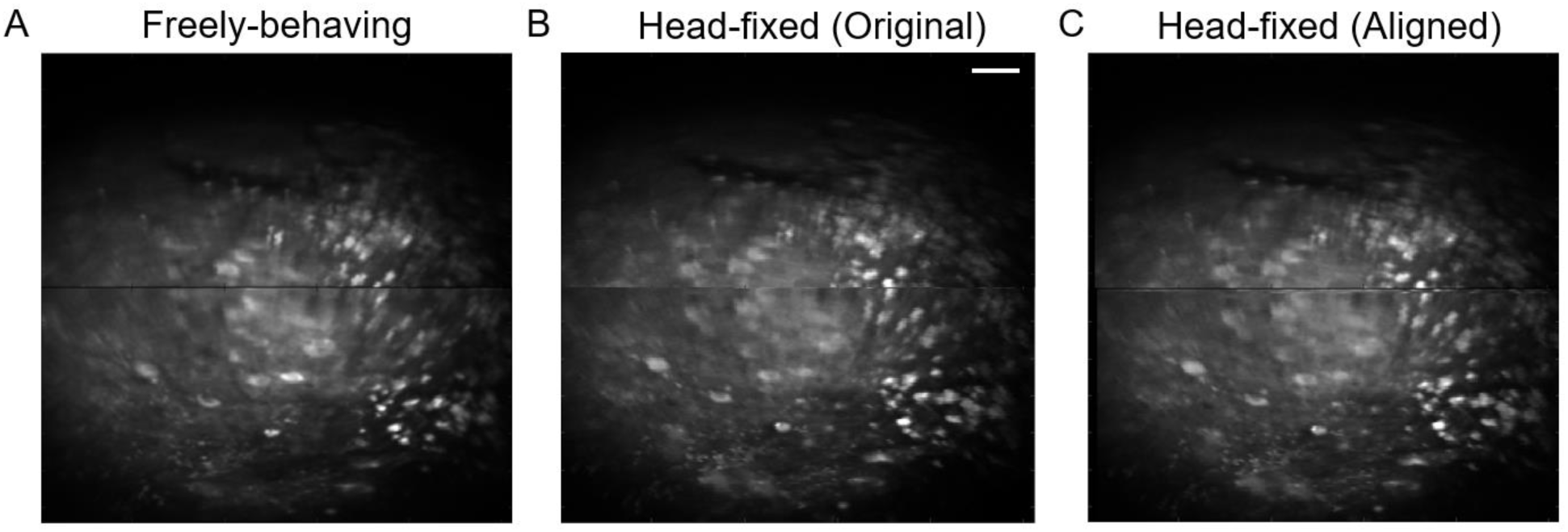
Field of view (FOV) comparison in mouse PFC between freely-behaving and head-fixed conditions. (A) Averaged intensity projection of calcium imaging in the mouse prefrontal cortex (PFC) during the freely-behaving condition. (B) Averaged intensity projection of the same region during the head-fixed condition. (C) Head-fixed imaging result spatially aligned to the result in the freely-behaving condition via translation shift based on maximum spatial correlation. Scale bar, (A-C), 50 μm.

**Figure S7.**
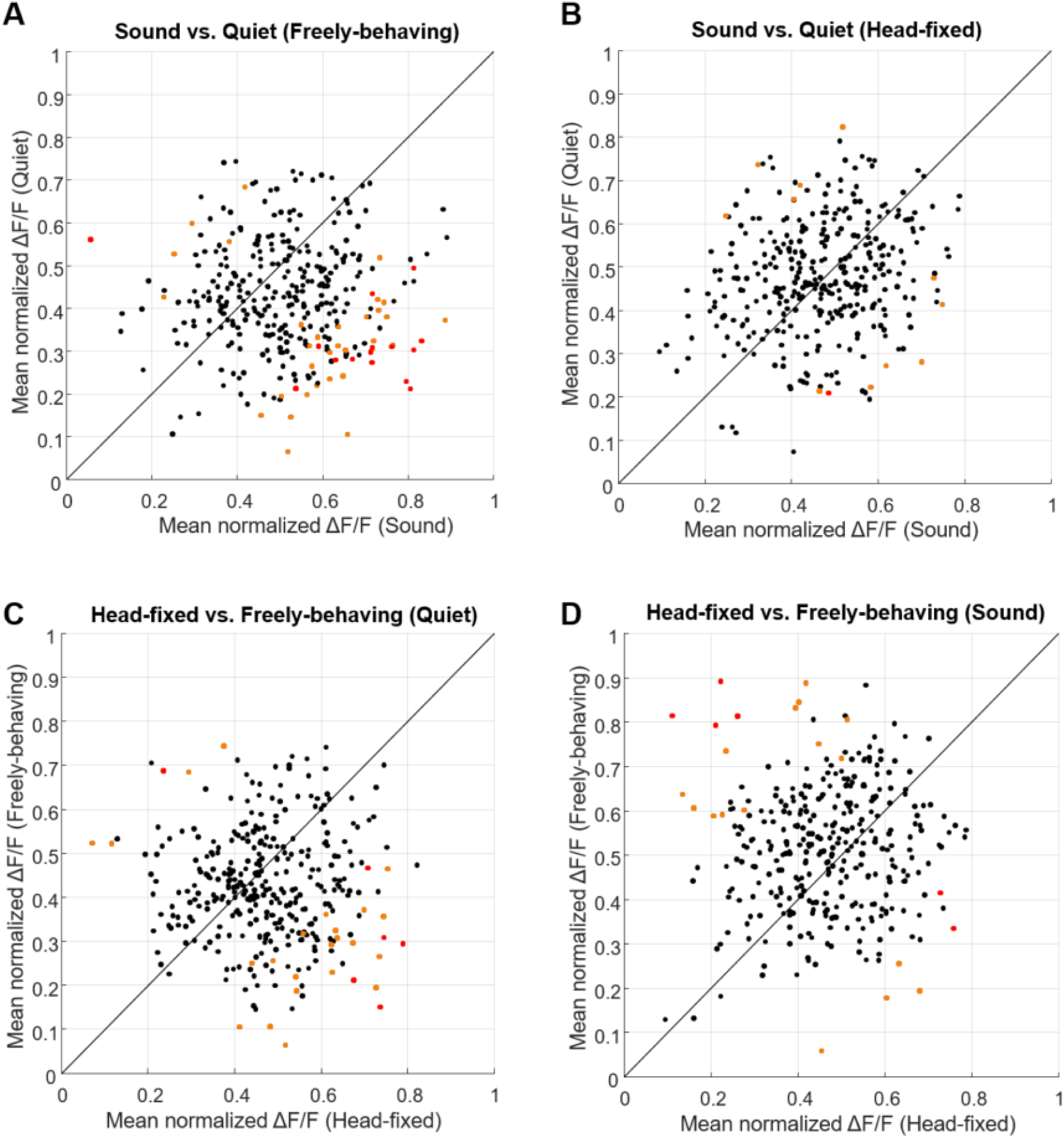
Neuronal response in mouse PFC in the auditory-stimuli experiments during head-fixed and freely-behaving conditions. Scatter plots compare mean normalized ΔF/F responses of neurons across different conditions. The mice were presented auditory stimulation with repeated sound-quite sessions, each lasting 45 seconds, in both freely-behaving and head-fixed conditions while maintaining the same imaging FOV. For each neuron, its activity level was normalized to 1 for the recordings in both freely-behaving condition and head-fixed condition. The data include 340 neurons from 2 mice. Each dot represents an individual neuron. The black diagonal line represents the unity line. Neurons with significant differences between the conditions in comparison are color-coded: orange for p <0.05 and red for p <0.01 (student t-test). (A) Sound vs. Quiet, under freely-behaving condition: mean normalized ΔF/F responses during auditory stimulation compared to quiet conditions in freely-behaving mice. Sound-preferring neurons (red and orange dots in the bottom right) show increased activity during auditory stimulation, while a subset of quiet-preferring neurons exhibit greater activity in quiet conditions (red and orange dots in the top left). (B) Sound vs. Quiet, under head-fixed condition: mean normalized ΔF/F responses during auditory stimulation compared to quiet conditions in head-fixed mice. (C) Head-fixed vs. Freely-behaving, under quiet condition: comparison of neuronal activity in quiet conditions between head-fixed and freely moving states. (D) Head-fixed vs. Freely-behaving, under auditory stimulation condition: comparison of neuronal activity during auditory stimulation between head-fixed and freely-behaving states. These results highlight that both the auditory stimulation and behavioral states (head-fixed vs. freely-behaving) can influence the neuronal activity in prefrontal cortex.

## Notes

### Competing Interest Statement

The authors have declared no competing interest.

